# Latitudinal dependence of stability trends in marine plankton in the Cenozoic

**DOI:** 10.64898/2026.02.11.705334

**Authors:** Maike L. Morrison, Adam Woodhouse, Anshuman Swain

## Abstract

The temporal stability and spatial heterogeneity of global marine ecosystems under changing climates reveal how biodiversity persists or collapses. However, the deep-time evolution of these phenomena remains poorly understood. We reconstructed the stability landscape of pelagic plankton from the Cretaceous-Paleogene extinction to the present. We find that the Cenozoic was a period of punctuated volatility with high turnover post-extinction and during the Late Neogene cooling. Spatially resolved analysis revealed latitude-dependent trends: equatorial regions stabilized over time, whereas polar communities, especially in the Southern Ocean, destabilized. Equatorial regions homogenized from initially high heterogeneity, whereas polar communities showed the opposite pattern, including a latitudinal seesaw of spatial heterogeneity over the past 30 million years. These findings illuminate temporal stability and spatial heterogeneity dynamics across geologic timescales.

## Main Text

The temporal stability and spatial heterogeneity of ecological communities are essential to their function and resilience (McCann, 2000). Temporal stability has been central to ecology and conservation biology, providing insights into species coexistence and community persistence (Ives and Carpenter, 2007). Spatial heterogeneity is likewise a core ecological idea, key to understanding the relationship between the biotic and abiotic factors shaping communities and to studying phenomena such as dispersal, competition, and environmental gradients (Londe et al., 2020).

While most research on temporal stability and spatial heterogeneity in ecological communities focuses on species richness, the relationship between taxonomic and functional composition is integral to how ecological communities maintain these properties under changing conditions. Functional redundancy, in which multiple species occupy similar ecological niches, enables the taxonomic composition of a community to fluctuate without disrupting its functional capacity (Fetzer et al., 2015). Conversely, systems may experience catastrophic functional loss with minimal species extinction if critical ecological roles are compromised (Säterberg et al. 2013; Fetzer et al., 2015). Thus, understanding temporal stability and spatial heterogeneity requires examining both taxonomic diversity and the functional roles that species perform within communities.

However, most studies of these phenomena draw on ecological data that are, on average, 3-5 years long and may not capture the variability within communities at longer timescales (Woodhouse et al., 2024). To characterize the spatial and temporal responses of ecological communities to the low-frequency, high-magnitude disturbances that have shaped Earth’s history, we need to incorporate long-term archives, such as the fossil record, but at a higher spatiotemporal fidelity than is commonly represented in paleontological studies (Woodhouse et al., 2024; Hull et al., 2015). Understanding these spatiotemporal dynamics requires integration of functional perspectives across eco-evolutionary time scales to comprehend how biological communities respond to abiotic forcing, including shifts in ocean circulation, temperature gradients, and carbon cycling.

The marine microfossil record, particularly that of planktonic foraminifera, provides a unique continuous archive for addressing questions of temporal stability and spatial heterogeneity over the last 66 million years of global change (Aze et al., 2011; Fraas et al., 2015; Swain and Woodhouse et al., 2024). Planktonic foraminifera serve as ideal model organisms for these questions due to their global distribution and exceptional preservation in the fossil record, making their record the largest fossil dataset available to science (Aze et al., 2011; Fenton & Woodhouse et al., 2021). More importantly, the functional ecology and morphology of other fossil groups are generally interpreted qualitatively, based on ecological and taxonomic uniformitarianism. Planktonic foraminifera, on the other hand, represent the only organismal group with a fully resolved phylogeny (including all the fossil species) spanning an entire geological era (Cenozoic, 66-0 Ma), within which the functional niche has been directly quantified via species-specific geochemical analyses (Aze et al., 2011; Fenton & Woodhouse et al., 2021; Woodhouse et al. 2023). This quantitative functional framework enables direct examination of how functional redundancy influences temporal stability and spatial heterogeneity across deep time.

In this work, we use the F_ST_-based Assessment of Variability across Abundances (FAVA) metric to quantify compositional variability in ecological communities across space and time. FAVA can be computed across a set of temporal samples, thereby serving as a proxy of the stability of the ecosystem, or across a set of spatial samples, in which case it serves as a measure of spatial heterogeneity. By joining the FAVA method with the Triton dataset (Fenton & Woodhouse et al., 2021), which provides detailed taxonomic, functional, and morphological data on foraminifera communities at unprecedented spatial and temporal scale, we reconstruct the temporal stability and spatial turnover of planktonic foraminiferal species and associated functional groups throughout the Cenozoic Era and across the globe (ecogroups and morphogroups; Woodhouse and Swain et al., 2023; Swain and Woodhouse et al., 2024). These analyses capture the spatial and temporal responses of foraminifera communities to major, low-frequency climatic events, such as hyperthermals and the greenhouse-to-icehouse transition, serving as a valuable resource for understanding ecological communities’ responses to major climatic shifts.

### FAVA measures temporal stability and spatial heterogeneity

The FAVA statistic is a measure of compositional variability across two or more compositional vectors, which here represent the ecogroup, morphogroup, or species composition of a foraminifera community sample at a particular time or place (e.g., vertical bars in Figure 1E-G). Computing FAVA across a set of time samples yields a measure of temporal stability of the foraminifera communities over the focal time period (temporal FAVA, tFAVA), whereas computing FAVA across spatial samples gives a measure of spatial heterogeneity (spatial FAVA, sFAVA). FAVA equals 0 if all the community samples have identical composition (high stability or homogeneity across samples). In contrast, FAVA equals 1 if each community sample is composed entirely of a single species, morphogroup, or ecogroup (high instability or heterogeneity). The mathematical properties of FAVA (Morrison et al., 2025) ensure the robustness of direct comparisons between datasets with different numbers of categories (e.g., FAVA values from ecogroups, morphogroups, or species). More detail on our FAVA analyses is provided in the Online Methods.

**Figure 1:**
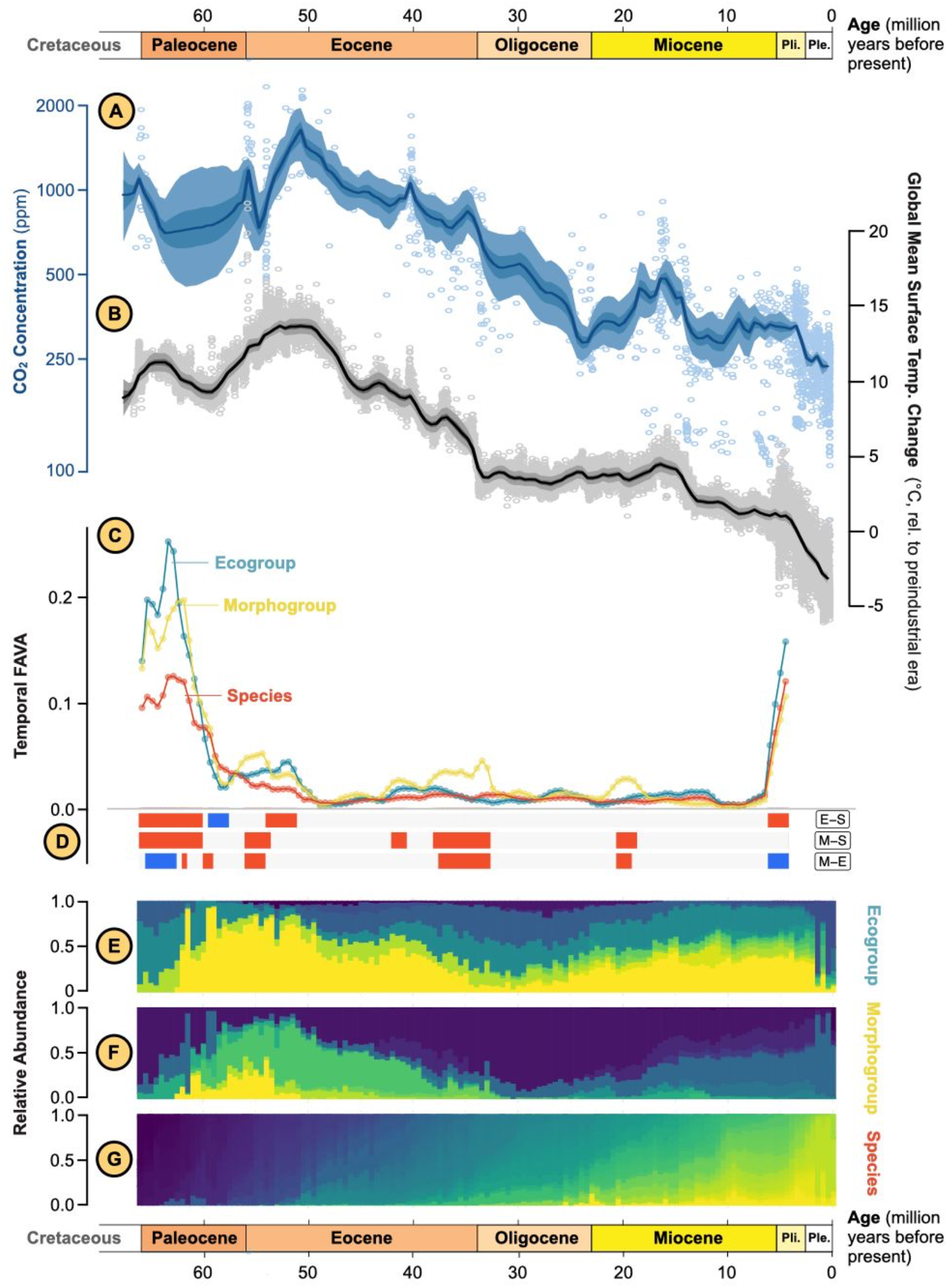
Overview of global climate variables (A-B), foraminifera composition (E-G), and temporal heterogeneity of foraminifera composition (C-D) over the past 66 million years. (A) CO_2_ concentration over the past 66 Myr. (B) Difference in global mean surface temperature relative to the pre-industrial era. Data in A-B from CenCO2PIP (2023). (C) Temporal FAVA values computed in sliding windows 10 samples (5 million years) wide for each data type. Figure S2 presents the horizontal span of each window, as well as results using alternative window widths. The x-value of each dot is the age of the oldest sample in the window; the y-value is temporal FAVA computed across all 10 samples in the window. (D) Colored bars along the bottom of the plot correspond to the difference in tFAVA values between ecogroups and species (E-S, top bar), morphogroups and species (M-S, middle bar), or morphogroups and ecogroups (M-E, bottom bar). Bars are red if the difference in tFAVA values is greater than 0.015, blue if the difference is less than -0.015, and gray otherwise. Many of these red or blue bars correspond to periods of significantly different tFAVA values between ecogroup and species (two-way Wilcoxon rank-sum tests, 5-million-year WB 66-60.5 Ma P=0.0001, 59.5-58 Ma P=0.114, 54-51.5 Ma P=0.002, 6-4.5 Ma P=0.083), morphogroup and species (two-way Wilcoxon rank-sum tests, 5-million-year WB 66-60.5 Ma P=0.0002, 56-54 Ma P=0.008, 42-41 Ma P=0.100, 38-33 Ma P=0.000003, 20.5-19 Ma P=0.029), or morphogroup and ecogroup (two-way Wilcoxon rank-sum tests, 5-million years beginning 65.5-63 Ma P=0.004, 62 Ma is 1 sample, 60-59.5 Ma P=0.333, 56-54.5 Ma P = 0.029, 37.5-33 Ma P=0.00001, 20.5-19.5 Ma P=0.100, 6-4.5 Ma P=0.343). (E-G) Temporal compositions of foraminifera at different taxonomic or functional data types: ecogroup (E), morphogroup (F), or species (G). Colors represent different categories within each data type. Each vertical bar represents the relative composition of all Triton samples within a 500,000-year window, regardless of sampling location.

### The Cenozoic is bookended by periods of dramatic instability

We first analyze the global foraminifera community, computing FAVA in 5-million-year-wide sliding windows to assess temporal stability of foraminifera over the Cenozoic. This application of FAVA to the Triton dataset unveils a Cenozoic history defined by distinct regimes of stability and instability (Figure 1C). We identify a U-shaped pattern of global instability, highlighting two periods of anomalously elevated instability: the recovery following the Cretaceous–Paleogene extinction and the recent bipolar cryosphere variability. This U-shaped pattern suggests that pelagic stability is not a static property of the marine realm but is highly sensitive to major transitions in Earth’s climate state.

By computing FAVA not only using species abundances, but also using abundances of ecogroups or morphogroups, we can disentangle patterns of functional redundancy and morphological disparity across the Cenozoic, linking these phenomena to major climatic events. Trends in temporal variability are highly correlated across the three data types, with all three exhibiting a distinct U-shaped pattern (Figure 1C). However, there are several periods of significant difference among the categories (Figure 1D). When the temporal variability of species significantly exceeds that of ecogroups or morphogroups, we say the community is exhibiting functional redundancy–the capacity for taxonomic categories to turn over while preserving functional composition. On the other hand, when the temporal variability of morphogroups or ecogroups significantly exceeds that of species, we say the community is exhibiting morphological or ecological disparity, which occurs when there is substantial morphological or ecological restructuring that is composed of many small changes in species abundances. Across the Cenozoic, we observe that functional redundancy is rare, occurring only briefly at the Paleocene/Eocene boundary and with tenuous statistical significance. Morphological disparity, on the other hand, occurs regularly throughout the Cenozoic.

### High instability during recovery from the K/Pg extinction

During the Greenhouse interval following the K/Pg extinction, the recovery of pelagic ecosystems was marked by a unique period in which both ecogroup and morphogroup instability significantly exceeded species-level variability (windows beginning 66-60.5 Ma, sample interval 66-56 Ma). This period of morphological and ecological disparity indicates a massive rearrangement of ecological and functional niches as the marine carbon pump and water column structure were restored. The rapid establishment of survivor morphogroups within the first few hundred thousand years was followed by a resurgence of oligotrophy, facilitating the rise of muricate clades like *Igorina, Morozovella*, and *Acarinina* (Aze et al. 2011). These clades, equipped with spines and algal symbionts, exploited novel upper-ocean niches, fundamentally restructuring the community beyond simple taxonomic replacement. As the Earth transitioned from the Paleocene oxygen isotope maximum toward the Early Eocene Climatic Optimum, a period of possible functional redundancy emerged (windows beginning 59.5-58 Ma, P=0.114). This suggests that the initial functional restructuring enabled a subsequent pulse of taxonomic diversification, where multiple species filled pre-existing ecological roles, providing a buffer of redundancy that increased community resilience.

### Recurring morphological instability following major climatic and evolutionary events

From the Eocene to the end of the Miocene, foraminiferal communities exhibited remarkable stability that was broadly consistent across functional and taxonomic classifications. However, there were three notable exceptions to this stability, all of which showed significantly elevated morphogroup tFAVA values relative to both species and ecogroup tFAVA. These periods occurred (1) at the onset of the Paleocene-Eocene Thermal Maximum (PETM) (beginning ∼56 Ma), (2) at the end of the Eocene (beginning ∼38 Ma), and (3) in the early Miocene (beginning ∼20.5 Ma).

Because each morphological category can contain multiple ecological and taxonomic categories, the foraminifera community can experience dramatic shifts in morphogroup composition, composed of many smaller shifts in ecogroup or species abundances, thereby generating much greater variability of morphogroups than of the other categories. When this occurs, the remaining morphogroups are populated by many relatively stable species, buffering the ecosystem against total functional collapse. This phenomenon also suggests that, during periods of substantial global change, it is morphogroups that are most responsive, suggesting their importance for monitoring the responses of foraminifera communities to future climatic perturbations.

(1) The onset of the Paleocene-Eocene Thermal Maximum (PETM) (∼56 Ma) was the most geologically rapid warming event of the Cenozoic (Kirtland Turner, 2018; Aze, 2022). Morphological malformation and short-lived morphologically unique excursion taxa have long been recorded from PETM sections (e.g., Bown & Pearson, 2009; Bralower et al., 2016), a possible explanation for the dramatically elevated morphogroup tFAVA during this period (Figure 1C-D). The continuation of high morphogroup tFAVA through the entirety of the EECO suggests that morphological disparity persisted well beyond the termination of the transient PETM (∼270 Kyr duration; Piedrahita et al., 2025).

Of note, during this period, it was not only morphogroups that experienced high tFAVA–ecogroups too saw a peak in FAVA, even exceeding morphogroup FAVA from 54-51.5 Ma (sample interval 54–47 Ma; P=0.002; Figure 1). Swain & Woodhouse et al. (2024) previously identified the post-PETM/EECO interval as a time of significant global ecological specialization amongst the planktonic foraminifera. As a whole, these results suggest that the transition to and from the stable Earth system states that bordered the unstable system of the EECO had significant repercussions for the planktonic foraminiferal phylogeny of the early Cenozoic, laying the groundwork for a period of relative stability that persisted for 9 million years—nearly the entire remainder of the Eocene.

(2) At the end of the Eocene, in the windows beginning 38 to 33 Ma (sample interval 38-28 Ma), morphogroup tFAVA again significantly exceeded tFAVA values of both species (P=0.000003) and ecogroups (P=0.00001). This decoupling of morphological instability from ecological and species instability aligns with global cooling and trophic restructuring following the Middle Eocene Climatic Optimum (MECO; 40.43-40.02 Ma), a protracted global warming event that interrupted the cooling trend initiated in the wake of the EECO (Henehan et al. 2020; Krause et al. 2023; Swain and Woodhouse, 2024). As temperatures dropped – as evidenced by the first record of ice-rafted debris in polar regions from ∼38 Ma (Eldrett et al. 2007) – the water column underwent significant physical changes (Flügel & Kiessling, 2002; Swain & Woodhouse et al., 2024). These shifts drove extinctions amongst key photosymbiotic planktonic foraminifera (38–34 Ma) (Lowery et al. 2020) that clustered across the Eocene-Oligocene Boundary (EOB; ∼33.9 Ma)--coinciding with a sharp peak in morphogroup tFAVA (Figure 1C). The EOB is typified by the initiation of permanent, continent-scale ice sheets across East Antarctica and is associated with global palaeoceanographic changes, leaving the oceans largely devoid of planktonic foraminiferal photosymbiosis until the mid-Miocene (Hutchinson et al. 2021; Boscolo-Galazzo et al. 2022). The EOB immediately precedes a rapid proliferation of the number of species within each morphogroup (Figure S1). The high redundancy within surviving morphogroups likely buffered the ecosystem against functional collapse, even as the late Eocene thermal and trophic changes facilitated major evolutionary events like the rise of crown cetaceans (∼39 Ma) (Figure S1; Marx & Fordyce, 2015).

(3) Finally, morphogroup tFAVA prominently exceeded both ecogroup and species tFAVA from 20.5-19 Ma (Figure 1C). This period is coincident with synchronous changes in global morphogroup richness and morphogroup community structure identified by Swain & Woodhouse et al. (2024), as well as significant shifts in marine food webs and nutrient cycling. For example, this interval includes a significant extinction amongst pelagic sharks (Sibert & Rubin, 2021) alongside a shift in foraminiferal-bound nitrogen isotopes suggestive of a shifting ocean oxygenation and/or nutrient cycling regime (Auderset et al., 2022). The abiotic mechanisms driving these changes across marine food webs are currently unknown, though efforts are underway to resolve them (Woodhouse & Kasbohm, 2026).

### Increasing instability descending into the bipolar icehouse

The dramatic rise in tFAVA values across all data types during the last ∼6 Myrs (Figure 1C) represents an evolutionary regime change within the planktonic foraminifera, which may elude the occurrence of a geologically recent regime shift like diversification across the Earth. From ∼7 Ma, the planet began to cool substantially as it switched from a unipolar to bipolar icehouse climate state (Herbert et al. 2016; Steinthorsdottir et al. 2021). Throughout these 7 Myrs of cooling, a series of evolutionary innovations took place amongst the planktonic foraminifera, which included: a wave of photosymbiont acquisition (Takagi et al. 2024), the rise and proliferation of a high number of extant cryptic species (Weiner et al. 2015; de Vargas et al. 2016), and a global reorganisation and equatorward migration of ecological communities (Woodhouse et al. 2023; Woodhouse & Swain et al. 2023; Swain & Woodhouse et al. 2024; Larina et al. 2025). Collectively, these events, alongside our tFAVA results, support the hypothesis of Woodhouse et al. (2023) that the compounded effects of seaway closures and steepening latitudinal and vertical marine temperature gradients, which typify the late Neogene, may have led to ecological and evolutionary innovations due to higher heterogeneity amongst marine ecological niches.

### Latitudinal gradient of stability trajectories

Up to this point, our analysis has focused on compositional dynamics of the global community, integrated over all spatial sampling locations. However, a spatially resolved analysis of these stability trends demonstrates that global averages obscure critical latitudinal dependencies. In general, tropical communities were most unstable (high tFAVA) immediately after the K/Pg extinction but have stabilized over time, whereas high-latitude communities, particularly those in the Southern Ocean, were relatively stable (low tFAVA) following the mass extinction but have become progressively more unstable toward the present day (Figure S3). We refer to these slopes of tFAVA over time as stability trajectories, and we find that they have a strong quadratic relationship with paleolatitude (Figure 2A, Table S1), a pattern we refer to as a latitudinal gradient of stability trajectories.

**Figure 2:**
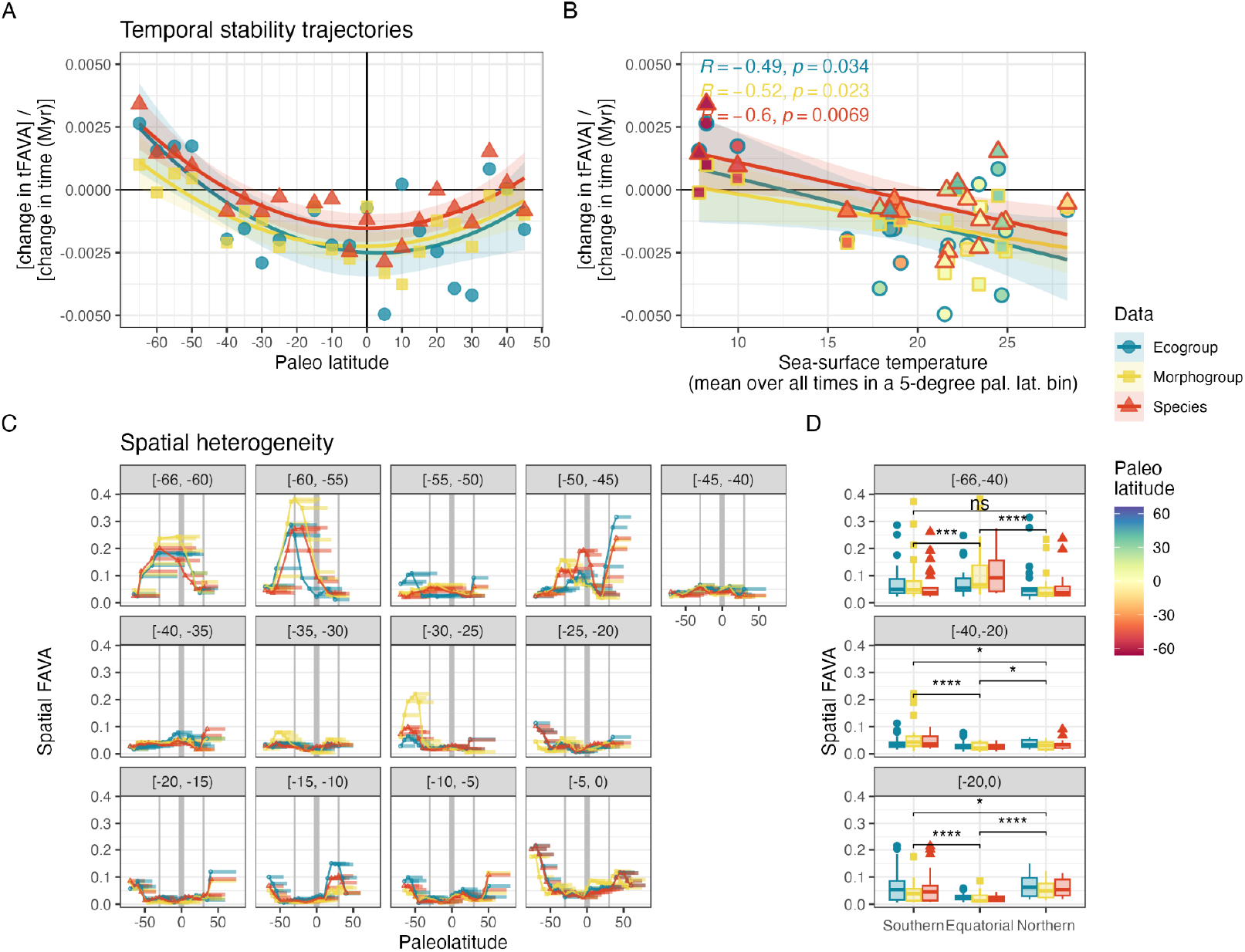
Latitudinal gradients of temporal stability trajectories (A-B) and spatial heterogeneity (C-D). (A) Each point represents the change in temporal FAVA (tFAVA) values (computed in sliding windows across four 5-million-year samples) over time (in millions of years) for one data type (color) in one paleolatitude bin. These slopes, which we refer to as stability trajectories, and the corresponding data they were computed on, are presented in Figure S3. Each point represents the southernmost bound of a 5-degree paleo latitude bin. Statistical significance of each slope is presented in Figure S4. Quadratic regression results are presented in Table S1. (B) Each point represents one paleo latitude bin: the x-axis presents the mean sea surface temperature in that bin (Figure S7B) and the y-axis presents the change in temporal FAVA over time in that bin (Panel A, Figure S7C). There is a significant, negative relationship between the change in tFAVA over time and sea surface temperature (inset R value presents Pearson correlation) for each of the three data types. Linear regression results are presented in Figure S7. (C) Samples are aggregated in bins 5 Ma wide (except the oldest bin, [-66, - 60) My) and 5 degrees paleolatitude wide (except the southernmost bin, (-76,70], and the northernmost bin, (80,88]). Spatial FAVA is computed in sliding windows 5 samples (approx. 25 degrees paleolatitude) wide within each temporal bin (panel). Horizontal bars present the value of sFAVA over the span represented by each sliding window. There is a dot at the southernmost bound of each sliding window, and lines connect these dots. Thick vertical grey bars denote the equator. Thin vertical grey bars denote +/-30 degrees paleolatitude. (D) sFAVA values from panel C are summarized according to temporal bin (panel), relationship to equator (x-axis), and data type (color). Each row of panel C is summarized in the equivalent row of D. “Equatorial” windows are contained within +/-30 degrees paleolatitude, whereas “southern” windows are south of these boundaries and “northern” windows are to the north. Asterisks represent two-sided P-values for Wilcoxon rank sum tests comparing sFAVA values between distinct equatorial regions, irrespective of data type. ns: P > 0.05; *: P <= 0.05; **: P <= 0.01; ***: P <= 0.001; ****: P <= 0.0001. P-values for each data type considered separately are presented in Figure S8. P-values at a more granular time resolution for the past 30 My are presented in Figure S10. Results for panels C and D are broadly consistent even under a much more conservative thresholding scheme (Figure S11).

The integration of these temporal stability trajectories (Figure 2A-B) with paleotemperature proxies reveals that these spatial patterns are fundamentally temperature-dependent. The significant negative correlation between stability trajectory and both δ^18^O (R=-0.74 for ecogroups, P=0.00013; Figure S6) and sea surface temperature (Figure 2B, S7) demonstrates that warmer regions – tropical latitudes—have stabilized over time, while cooler regions – polar latitudes – have destabilized. This relationship is strongest for ecogroups, suggesting ecological categorizations are most responsive to thermal forcing and that the observed latitudinal gradient in stability trajectories reflects the differential climatic histories experienced across latitudes during the Cenozoic cooling trend

Collectively, these results indicate that the current stability observed in tropical pelagic ecosystems is a geologically recent phenomenon. In contrast, modern polar-region volatility is driven by the late Neogene transition to a bipolar cryosphere (Woodhouse and Swain et al., 2023). We demonstrate that these stability trajectories are not correlated with sampling effort (Figure S5) or with our statistical confidence in the slopes (Figure S4). Ultimately, the U-shaped global trajectory of temporal instability over the Cenozoic (Figure 1C) is an aggregate of these divergent latitudinal processes (Figure 2A). The stabilization of the tropics and the destabilization of the poles are two sides of the same macroevolutionary coin, driven by the steepening of latitudinal temperature gradients. As the planet continues to experience rapid environmental change, these deep-time insights suggest that polar ecosystems are currently in a state of geologically anomalous instability, whereas the historically stable tropical refugia may face new thresholds of functional restructuring if modern warming reverses the stabilization trends of the last 60 million years.

### Reorganization of spatial heterogeneity

To this point, our analyses have used FAVA to assess the temporal stability of foraminiferal communities by quantifying compositional variability across temporal samples. Now, we pivot to using FAVA as a measure of the spatial heterogeneity of these communities, quantifying compositional variability across samples representing different paleolatitude bins. In this context, FAVA can be considered a measure of ecological beta diversity. We find that in the early Cenozoic, the tropics exhibited the highest spatial heterogeneity, reflecting the patchy and localized nature of ecosystem recovery following the mass extinction (Figure 2C, S8). However, throughout the Cenozoic cooling, equatorial regions have grown increasingly spatially homogeneous (Figure S9). By the late Neogene, the locus of spatial turnover had migrated to the poles. The Southern Ocean in particular now exhibits intense spatial heterogeneity, mirroring the high temporal instability observed in high-latitude assemblages during this period. This reversal from an equator-dominated spatial heterogeneity regime to a pole-dominated one highlights the major impacts of global cooling on the spatial organization of marine biodiversity and likely contributed to the latitudinal gradient in temporal stability trajectories documented in Figure 2A.

During the early Cenozoic warmhouse interval (66-55 Ma; Westerhold et al. 2020), communities exhibited elevated spatial heterogeneity in low-latitude regions, particularly south of the equator (Figure 2C), reflecting the spatially heterogeneous restructuring of ecological niches following the K/Pg extinction. Tropical regions experienced the most dramatic reorganization, with evidence for the initial requisition of photosymbiosis in planktonic foraminifera occurring in the southern hemisphere (Quillévéré et al. 2001; Swain & Woodhouse et al. 2024). The subsequent hothouse interval (55-50 Ma; Westerhold et al. 2020) marked an unexpected transition, during which morphogroup and species spatial FAVA were very similar, whereas ecogroup spatial FAVA values diverged, showing elevated values at high southern latitudes. A similar observation was noted by Swain & Woodhouse et al. (2024), suggesting the existence of ecological refugia in high latitudes due to thermally prohibitive low-latitude global temperatures during the hothouse.

The Warmhouse (45-30 Ma) and Coolhouse (30-5 Ma) intervals exhibited relative spatial homogeneity across latitudes, consistent with the prolonged mid-Cenozoic stability evident in temporal analyses (Figure 1C). Significantly, however, the past 30 Myrs exhibit a “latitudinal seesaw” in spatial heterogeneity dynamics (Figure 2C, S10). High-southern-latitude foraminifera communities showed the highest spatial heterogeneity from 30-25 Ma, followed by spatially heterogeneous northern communities from 15-10 Ma, and concluding with both northern and southern communities experiencing greater heterogeneity than equatorial communities during the past 5 Myrs (Figure S10). The initiation of this signal in the early Oligocene (30-25 Ma) suggests that early Antarctic glaciations may have driven high morphogroup community turnover in close proximity to the newly established ice sheet, possibly in response to the dynamic waxing and waning of Antarctic ice extent and associated regional palaeoceanographic changes (Liebrand et al. 2016; Swain & Woodhouse et al. 2024). The northern hemisphere peak, which reached prominence from 15-10 Ma, was situated in the low to mid-latitudes (Figs 2c, S11a). This approximates the time where global plankton populations and biogeochemical modelling suggest global cooling following the Miocene Climatic Optimum (17-14 Ma; Figure 1a) facilitated the expansion of organisms into the oceanic twilight zone (Boscolo-Galazzo & Crichton et al. 2021; Boscolo-Galazzo et al. 2022; Woodhouse & Swain et al. 2023). Why this phenomenon was isolated to the northern hemisphere from 15-10 Ma is unclear; however, model and proxy data of the latitudinal temperature gradient consistently suggest that it asymmetrically steepened towards the southern hemisphere through the mid-Miocene due to Antarctic glaciation (You et al. 2009; Pound et al. 2012). The latitudinal seesaw in sFAVA values from south to north may therefore reflect the asynchronous surpassing of a temperature threshold by planktonic foraminifera in one hemisphere relative to the other as the Earth system cooled asymmetrically through 30-10 Ma.

The modern icehouse (5-0 Ma) established a pattern of poleward-dominated volatility, with intense spatial heterogeneity in the Southern Ocean and, to a lesser extent, northern high latitudes (Figure 2C, S10). This pattern correlates strongly with the temporal destabilization of high-latitude assemblages documented in Figure 2D, further supporting the interpretation that polar instability represents a geologically recent phenomenon (Figure 1). The establishment of the northern hemisphere ice sheet and the advent of the bipolar cryosphere further strengthened global latitudinal temperature gradients and triggered substantial paleoceanographic reorganization (Lawrence et al. 2006; Rahaman et al. 2025). Indeed, modern polar volatility is likely a product of intensified oceanographic circulation, and the expansion of seasonal ice cover, all of which have created a higher degree of niche heterogeneity across latitudinal belts (Woodhouse et al. 2023; Woodhouse & Swain et al. 2023; Larina et al. 2025).

## Conclusions

In this paper, we analyze the spatiotemporal evolution of stability in Cenozoic planktonic foraminifera (66–0 Ma) using the FAVA metric. We find that a global U-shaped trajectory of instability–high during the post-extinction recovery and the modern icehouse–masks an important latitudinal divergence. Stability has evolved continuously in space and time: tropical communities have become progressively more stable and homogeneous, whereas high-latitude assemblages have steadily destabilized. Consequently, the planetary ‘locus of instability’ has migrated from the equator in the early Cenozoic Greenhouse to the poles in the modern Icehouse. This inversion correlates strongly with cooling temperatures, suggesting that the spatial distribution of pelagic stability is dynamically coupled to the Earth system’s thermal state. This work contributes a framework for studying the long-term turnover of species and functional abundances over space and time that does not focus on changes in richness, but instead captures the dynamics of the full community composition.

## Data and Code Availability

All code needed to replicate these analyses is available at https://github.com/MaikeMorrison/foram-fava. The data used in this manuscript are from the Triton database (see Fenton and Woodhouse et al., 2021). The statistical methods used in this manuscript are implemented in the R package “FAVA” which is available on CRAN.

## Acknowledgements

The authors would like to thank the Santa Fe Institute for facilitating this collaboration and Chris Kempes for helpful conversations.

