## Supplementary Materials for "Latitudinal dependence of stability trends in marine plankton in the Cenozoic"

### Online Methods

##### Data

We sourced occurrence data from the Triton dataset (Fenton & Woodhouse et al., 2021). We excluded microperforate and medioperforate wall textures (38,046 entries) due to insufficient phylogenetic resolution (Aze et al., 2011). For the remaining macroperforate species, speciation and extinction datums were applied in accordance with Aze et al. (2011) and Fenton & Woodhouse et al. (2021). Occurrences outside these stratigraphic ranges were trimmed to eliminate artificial "tails" attributable to reworking or identification error, resulting in a final dataset of 359,253 entries. Data were binned temporally at 0.5 Myr intervals and spatially in 5° palaeolatitudinal bands. Bins with total abundance less than 5 were not included. Species were assigned to the "ecogroups" and "morphogroups" of Aze et al. to determine ecological and morphological influences on community dynamics.

Ecogroups were assigned based on biogeography and shell stable isotopes (oxygen (δ18O) and carbon (δ13C)), which serve as proxies for bathymetric position and ecological niche (CITE). The ecogroups are: 1) symbiotic mixed layer; 2) asymbiotic mixed layer; 3) thermocline; 4) sub-thermocline; 5) high-latitude; and 6) high productivity/upwelling dwellers (Figure 1). Many species are assigned to two, three, or four ecogroups, and each of these combinations is treated as a distinct ecogroup category. Morphogroups consist of 19 forms defined by functional shell characteristics. Combined, these groups allow us to infer water column vertical structure and habitability through geological time (Woodhouse & Swain et al., 2023). Although the sequential evolution of these functional groups is well-documented, the Triton dataset facilitates a novel analysis of this group's response to environmental change at a higher spatiotemporal resolution.

##### FAVA method

We use FAVA, an $F_{ST}$-based Assessment of Variability across vectors of relative Abundances (Morrison et al. 2025), to measure the stability of planktonic foraminifera communities over time. In general, FAVA is a statistic that quantifies the compositional variability across two or more communities, equalling 0 when all the communities have identical composition and equalling 1 when each is composed entirely of a single category (Figure M1C). When computed across a set of longitudinal samples, FAVA can be thought of as a proxy for the temporal stability of the community, equalling 0 when the community has constant composition over time and equalling 1 when each time point is composed entirely of one species–necessitating large-magnitude fluctuations in community composition (Morrison et al. 2025). When computed across a set of spatial samples, FAVA can be thought of as a measure of the spatial heterogeneity of the community, equalling 0 when the samples are totally homogeneous in their composition, and 1 when there is complete spatial turnover from one sample to the next. Visually, FAVA increases as the stacked bar plot of relative abundances over time gets more jagged and uneven (Figure M1C).

Mathematically, FAVA is defined as a normalized difference of hierarchical computations of the Gini-Simpson index (Figure M1B), a measure of the diversity of a single community (see Morrison et al. 2025 for mathematical details). It is important to note that FAVA, which measures compositional variability across two or more communities, is complementary to and fundamentally different from diversity measures such as the Gini-Simpson index, which measure the diversity of individual communities (Figure M1A).

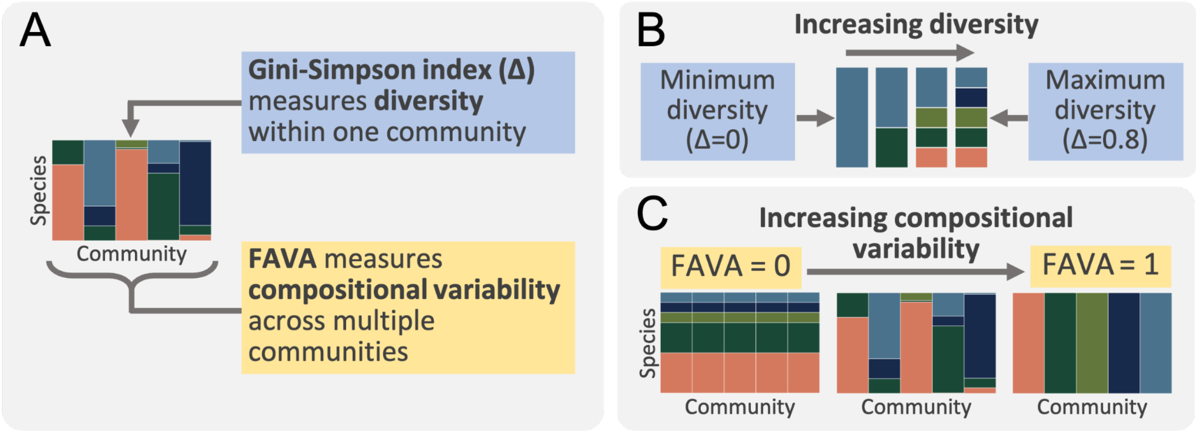


**Figure M1. Definitions of diversity and compositional variability.** Each vertical bar represents the composition of one foraminifera community (e.g., one timepoint), with colors corresponding to different species. Note that here "species" is used in place of any category (e.g., species, ecogroups, or morphogroups. Note also that when FAVA is unweighted (as it is here), FAVA and $F_{ST}$ have the same minimizing and maximizing cases. **(A)** Diversity and compositional variability (FAVA) measure fundamentally different quantities. **(B)** Diversity measures such as the Gini-Simpson diversity are minimized when a community is comprised entirely of a single species, and maximized when all species are evenly abundant (the maximum Gini-Simpson diversity for five species is 0.8; this quantity approaches 1 as the number of species increases). **(C)** FAVA equals 0 when all communities have identical composition, and 1 when each community is composed entirely of a single category.

We performed these analyses using the R package *FAVA*, which was downloaded from CRAN. We used built-in functions to weight FAVA by the time represented by each community composition vector, thereby accounting for any instances of uneven longitudinal sampling. Further, convenient mathematical properties of FAVA guarantee that it can be directly compared between data sets with different numbers of categories–allowing us to compare the FAVA values between foraminifera ecogroups, morphogroups, or species (Morrison et al. 2025).

**FAVA data analysis**

We present three FAVA-based analyses of foraminifera data. First, we analyzed the stability of foraminifera ecogroups, morphogroups, and species worldwide over the past 66 million years (Figure 1E-G). Each vertical bar in Figure 1E-G, which we call a "sample," represents the composition of the global foraminifera community during a period of 500,000 years. Dated foraminifera observations are assigned to these samples by rounding their dates to the nearest 0.5 My. For example, the sample labeled "55 My" includes observations between 54.75 and 55.25 My. Computing FAVA across a set of two or more adjacent time samples yields a measure of temporal stability of foraminifera composition during the period spanned by the selected samples. High temporal FAVA values reflect high compositional variability across the samples, whereas low temporal FAVA values reflect stability of composition across samples.

We computed temporal FAVA in sliding windows 10 samples (5 million years) wide in order to understand how the stability of the global foraminifera community depended on both time and data type (Figure 1C). These sliding window analyses follow Figure 4 of Morrison et al. (2025). In general, we refer to sliding windows by the time of the oldest sample in the window. Consider, for example, the sliding window beginning 55 Ma. This window includes the following 10 samples: (55, 54.5, 54, 53.5, 53, 52.5, 52, 51.5, 51, 50.5). From this list, it may seem that this window spans only 4.5 My. However, because each sample is defined by rounding the actual, unbinned foraminifera observation dates to the nearest 0.5, the full range of the actual, unbinned foraminifera observation dates included in this window is 55.25 to 50.25 Ma, which spans 5 My.

Second, we analyzed the stability of spatially-resolved foraminifera communities over the past 66 million years (Figure S3). In this analysis, each sample (vertical bars in Figure S3A) represents the composition of the foraminifera community in a paleolatitude bin 5-degrees wide during a period of 5 million years. Paleolatitude values range from -65 degrees to 49 degrees. We computed tFAVA in sliding windows 4 samples (20 million years) wide within each 5-degree paleolatitude bin in order to understand how the stability of the foraminifera community depends on time, data type, and paleolatitude (Figure S3B).

Third, we used FAVA to analyze the spatial heterogeneity of foraminifera communities, rather than their temporal stability. We bin samples into latitudinal bins 5-degrees wide, and into time bins 5 My wide (with the exception of the oldest bin, which is 6 My wide). We compute spatial FAVA in sliding windows 5-samples wide across paleolatitude bins–so each sliding window computation of FAVA spans approximately 25 My. The results of this analysis are presented in Figure 2C.

**Secondary analyses of FAVA results**

We performed a number of additional analyses on our temporal FAVA results in particular. We used ordinary least squares (OLS) linear regression models to measure the relationship between time and FAVA at each paleolatitude for each data type (Figure S3B). To perform these regression analyses, each sliding window was represented as 4 data points with x-values corresponding to each of the 4 sampling times included in the window and y-values corresponding to the single FAVA value computed for that window. For these regressions, a negative slope suggests that FAVA values are decreasing and therefore the community is growing more stable over time, whereas a positive slope suggests that FAVA values are increasing and consequently the community is growing less stable over time. We refer to these slopes as the stability trajectories for each paleolatitude bin. We used OLS quadratic regression models to quantify the relationship between paleolatitude and stability trajectory for each data type (Figure 2, Table S1).

In a separate analysis, we incorporated data from Gaskell et al. (2022) to understand the relationship between stability trajectories and a temperature proxy, using either $\delta^{18}O$ (Figure S6) or inferred sea-surface temperature (SST, Figure S7). $\delta^{18}O$ values were reported in 5 My intervals for the full range of paleolatitudes considered in our FAVA analyses. We binned these paleolatitude values using the same binning scheme used in Figure S3, computed the mean $\delta^{18}O$ value over time within each paleolatitude bin, and then used OLS linear regression models to quantify the relationship between stability trajectory and $\delta^{18}O$ (Figure S6). SST measurements were reported at the resolution of individual temporal and latitudinal samples. For consistency with the $\delta^{18}O$ analysis and to control for temporally uneven sampling effort, we first computed the mean SST in bins of 5 degrees paleolatitude and 5 million years and then computed the mean temperature across all time bins within each paleolatitude bin. We then used OLS linear regression models to quantify the relationship between stability trajectory and SST (Figure S7).

### Supplementary figures

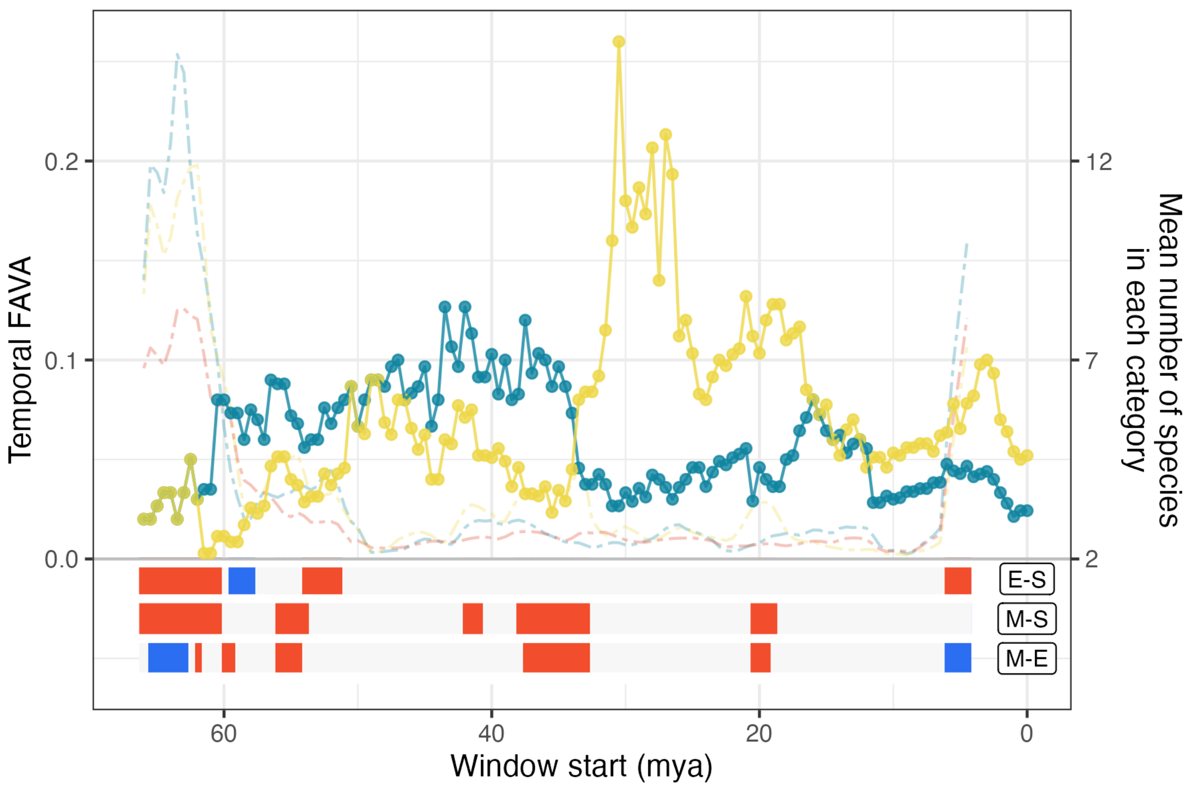


**Figure S1: Mean number of species within ecogroups (blue solid lines) or morphogroups (yellow solid lines) over time (refer to right y-axis).** Background (left y-axis) reproduces Figure 1B.

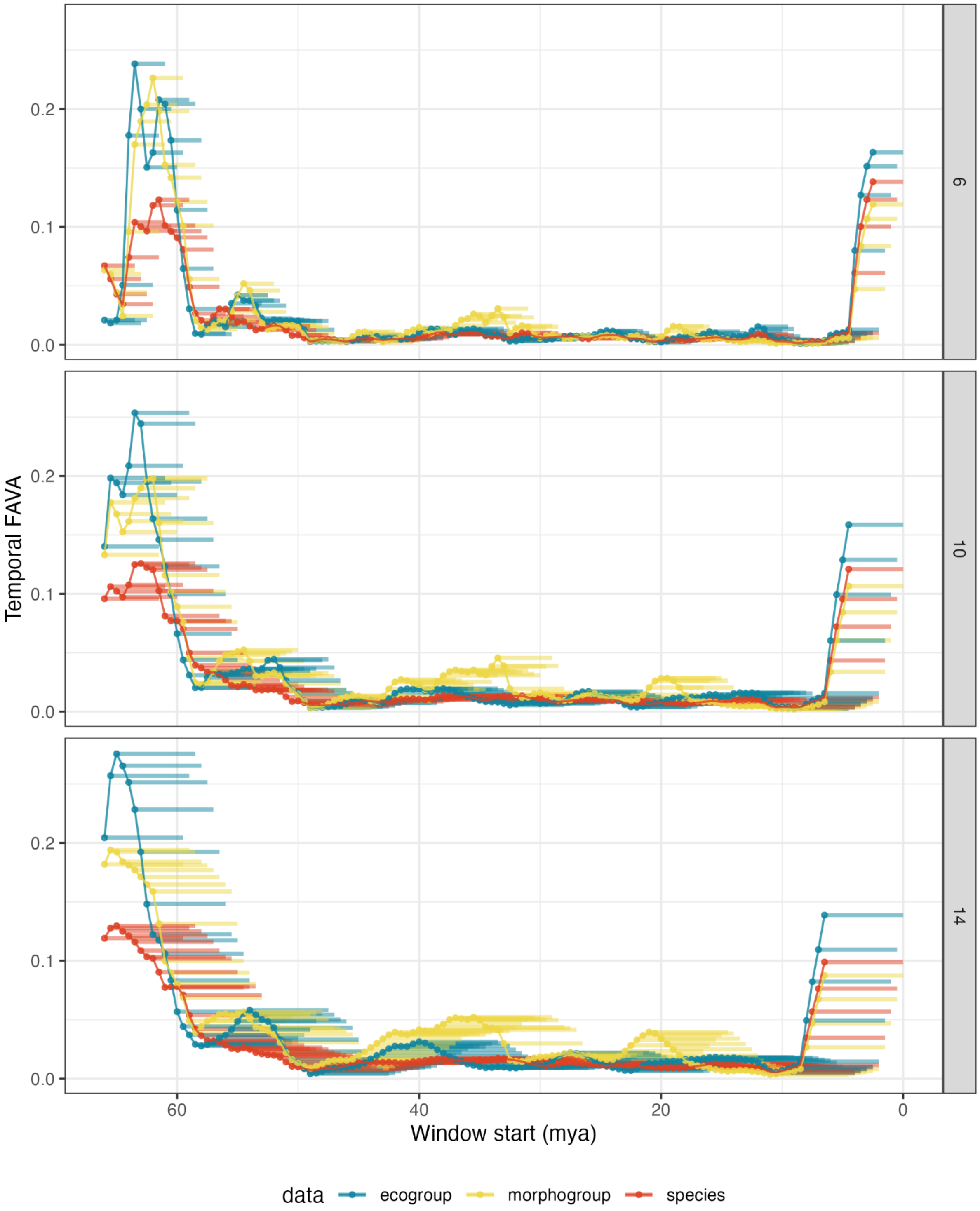


**Figure S2. Reproduces Figure 1b, varying the number of samples included in each sliding window.** Each row corresponds to the number of samples included in each sliding window (6, 10, or 14). The middle row (10 samples) reproduces Figure 1B. As in Figure 1B, points represent the oldest sample in each sliding window, and lines connect subsequent points. Here, the breadth of each sliding window is represented by a horizontal bar.

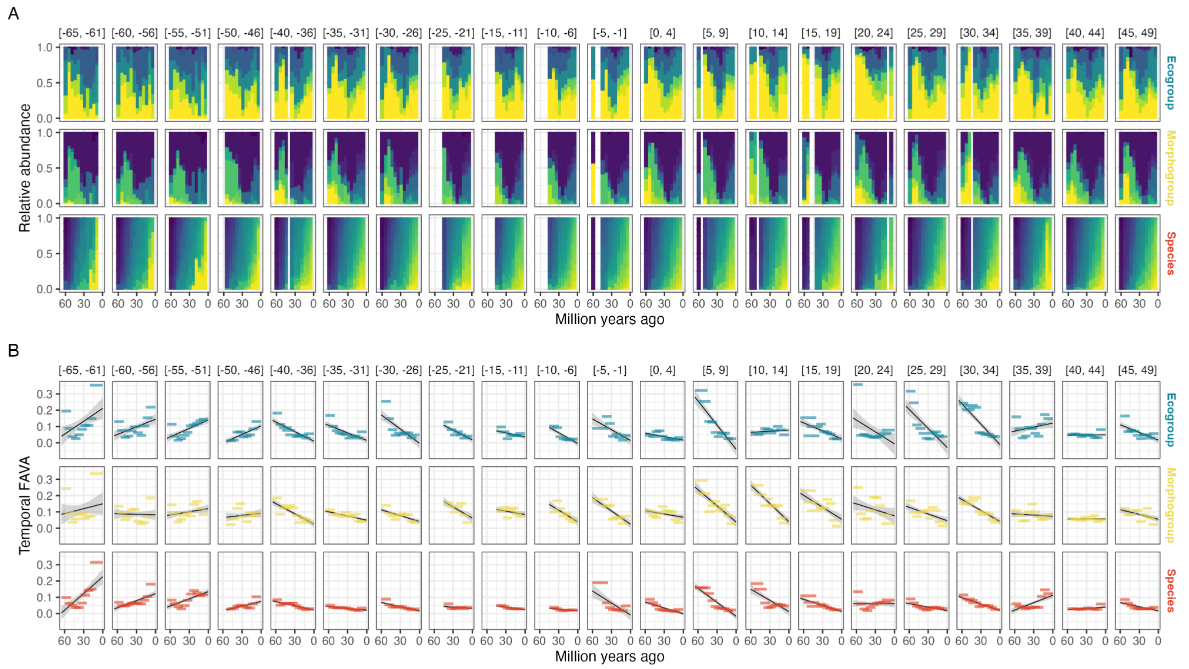


**Figure S3: Foraminifera composition (A) and stability trajectories (B) by latitude (columns) for each data type (rows).** In both panels, the x-axis represents time in millions of years ago, with the past on the left and the present on the right. (A) Each panel presents the relative abundances of categories of one foraminifera data type over time for each 5-degree bin of paleo latitudes. Within a panel, each vertical bar represents the foraminifera composition in a bin of sampling times 5 million years wide. Colors align with the colors in Figure 1a. (B) Temporal FAVA values computed in sliding windows 4 samples (20 million years) wide across the relative abundances samples presented in panel B. Black lines present linear slopes fit to the data, which are summarized in Figure 2A.

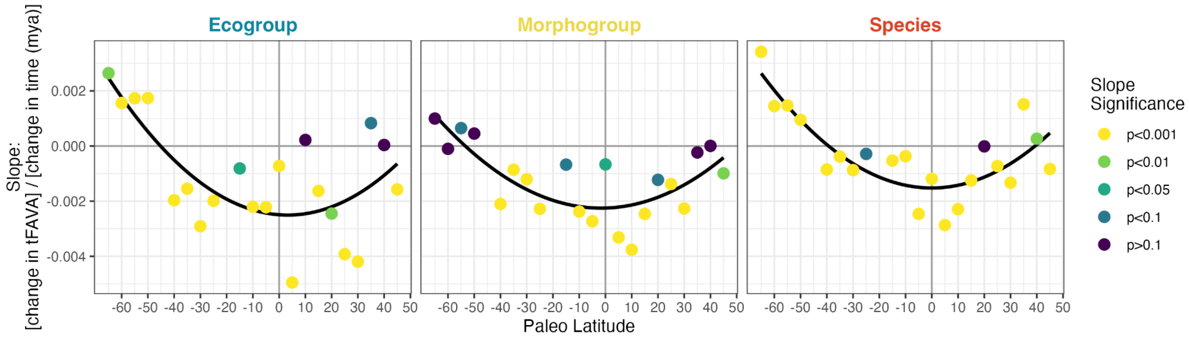


**Figure S4: Reproduces Figure 2A, with point colors corresponding to the significance of the corresponding slope, and each data type presented separately.**

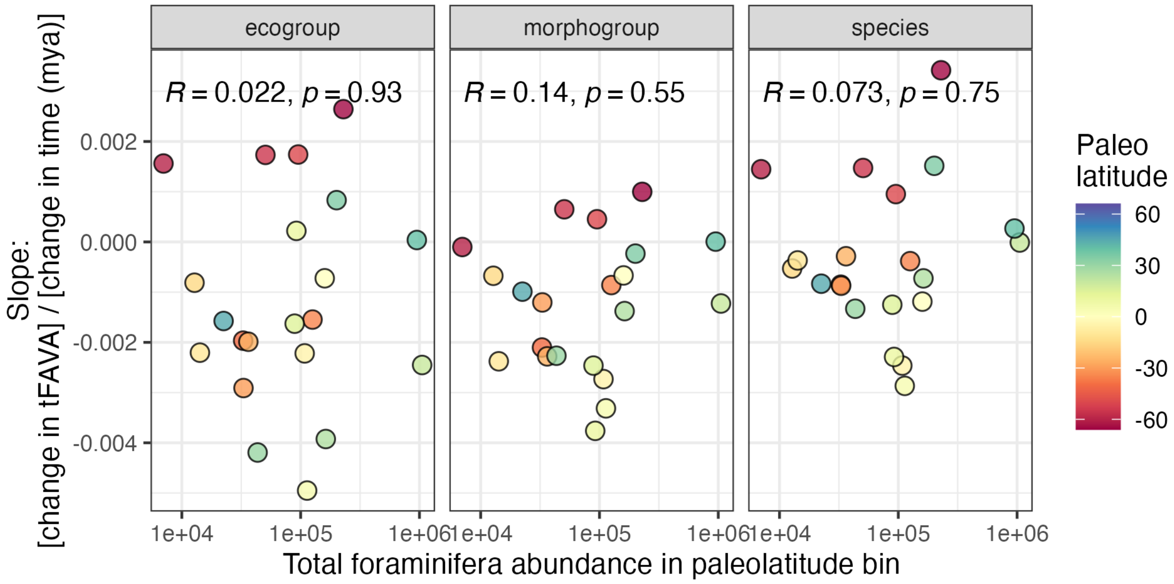


**Figure S5. Stability trajectories are not correlated with sampling effort.** Each point corresponds to one point in Figure 2A. Color represents the southernmost bound of a 5-degree paleo latitude bin, as in Figures S6 and S7. X-axis presents the total pseudoabundance of foraminifera sampled within each paleolatitude bin. Y-axis shows the stability trajectory, or change in temporal FAVA over time, within each paleolatitude bin, a proxy for sampling effort. Across the three data types, we see no significant Pearson correlation (*R*) between sampling effort and stability trajectory, suggesting that our observed latitudinal gradient is not driven by latitudinal differences in sampling.

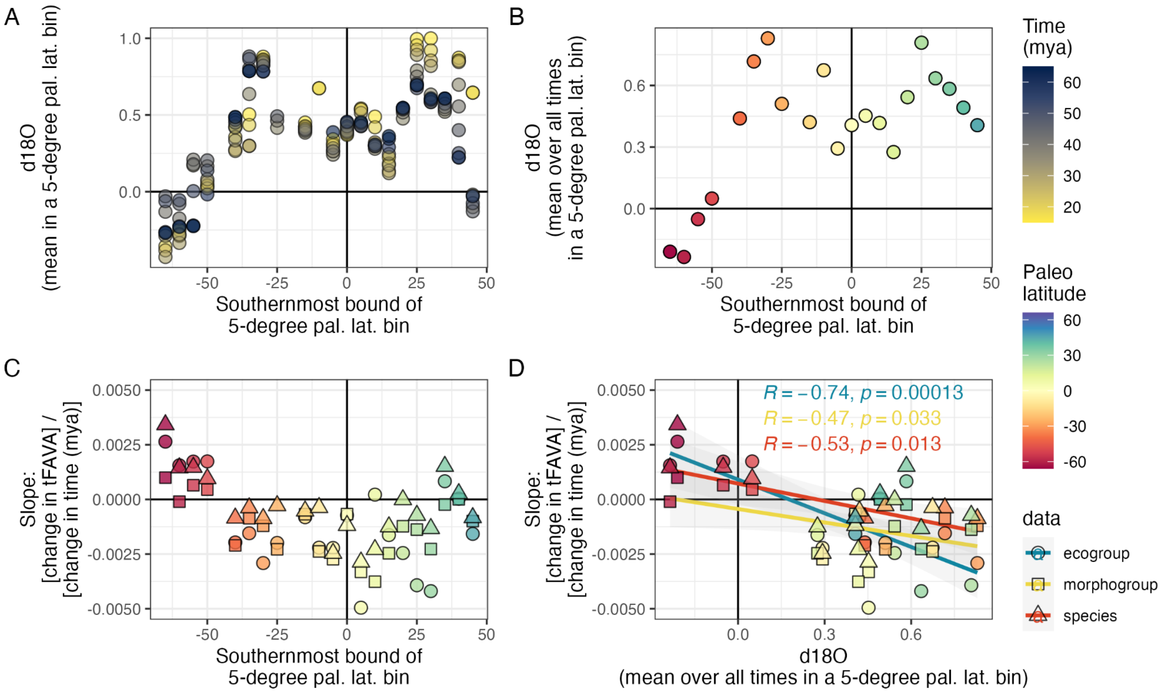


**Figure S6. Stability trajectories are correlated with integrated paleo temperature, as measured by mean** $\delta^{18}O$**.** $\delta^{18}O$ data is from Gaskell et al. (2022); we restrict the data to values in the past 65 million years. (A) $\delta^{18}O$ values are averaged within the same 5-degree paleo latitude bins used in Figures 2 and S3. x-axis presents the southernmost bound of each bin, y-axis presents the mean $\delta^{18}O$ in the bin, and color represents the time in millions of years ago for which that temperature applies. (B) Replicates panel A, except $\delta^{18}O$ values are averaged across years for each paleo latitude bin. We refer to these values as “integrated paleo temperature.” (C) Replicates Figure 2A, except data type is represented by shape and color represents the paleo latitude. (D) Each point represents one paleo latitude bin: the x-axis presents the mean $\delta^{18}O$ value in that bin (panel B) and the y-axis presents the change in tFAVA over time in that bin (panel C). There is a significant, negative relationship between the change in tFAVA over time and $\delta^{18}O$ (inset R value is Pearson correlation). Correlation between slope and $\delta^{18}O$ is strongest for ecogroups. Linear regression results are as follows.

Ecogroup: intercept = 0.0009 (P=0.104), slope = -0.005 (P=0.0001), adj. R^2 = 0.524.

Morphogroup: intercept = -0.0004 (P=0.33), slope = -0.002 (P=0.033), adj. R^2 = 0.176.

Species: intercept = 0.0007 (P=0.148), slope = -0.003 (P=0.013), adj. R^2 = 0.246.

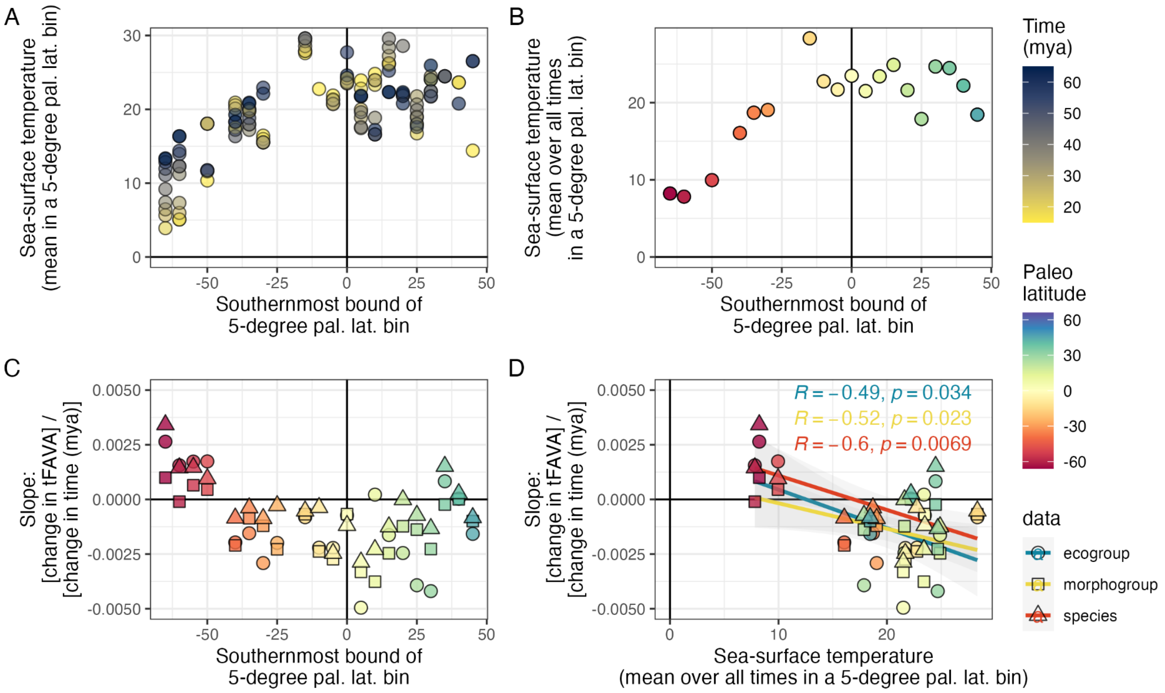


**Figure S7.Stability trajectories are correlated with integrated paleo temperature, as measured by inferred sea-surface temperature.** Sea-surface temperature (SST) is from Gaskell et al. (2022); we restrict the data to values in the past 65 million years. Note that we present both δ18O and SST because the data for δ18O is more complete. (A) SST values are averaged within the same 5-degree paleo latitude bins used in Figures 2 and S3. x-axis presents the southernmost bound of each bin, y-axis presents the mean SST in the bin, and color represents the time in millions of years ago for which that temperature applies. (B) Replicates panel A, except SST values are averaged across years for each paleo latitude bin. We refer to these values as “integrated paleo temperature.” (C) Replicates Figure 2A, except data type is represented by shape and color represents the paleo latitude. (D) Each point represents one paleo latitude bin: the x-axis presents the mean SST value in that bin (panel B) and the y-axis presents the change in tFAVA over time in that bin (panel C). There is a significant, negative relationship between the change in tFAVA over time and SST (inset R value is Pearson correlation) for each of the three data types. Linear regression results are as follows.

Ecogroup: intercept = 0.0022 (P=0.177), slope = -0.0002 (P=0.034), adj. R^2 = 0.193.

Morphogroup: intercept = -0.0001 (P=0.315), slope = -0.0001 (P=0.023), adj. R^2 = 0.225.

Species: intercept = 0.0026 (P=0.021), slope = -0.0002 (P=0.007), adj. R^2 = 0.319.

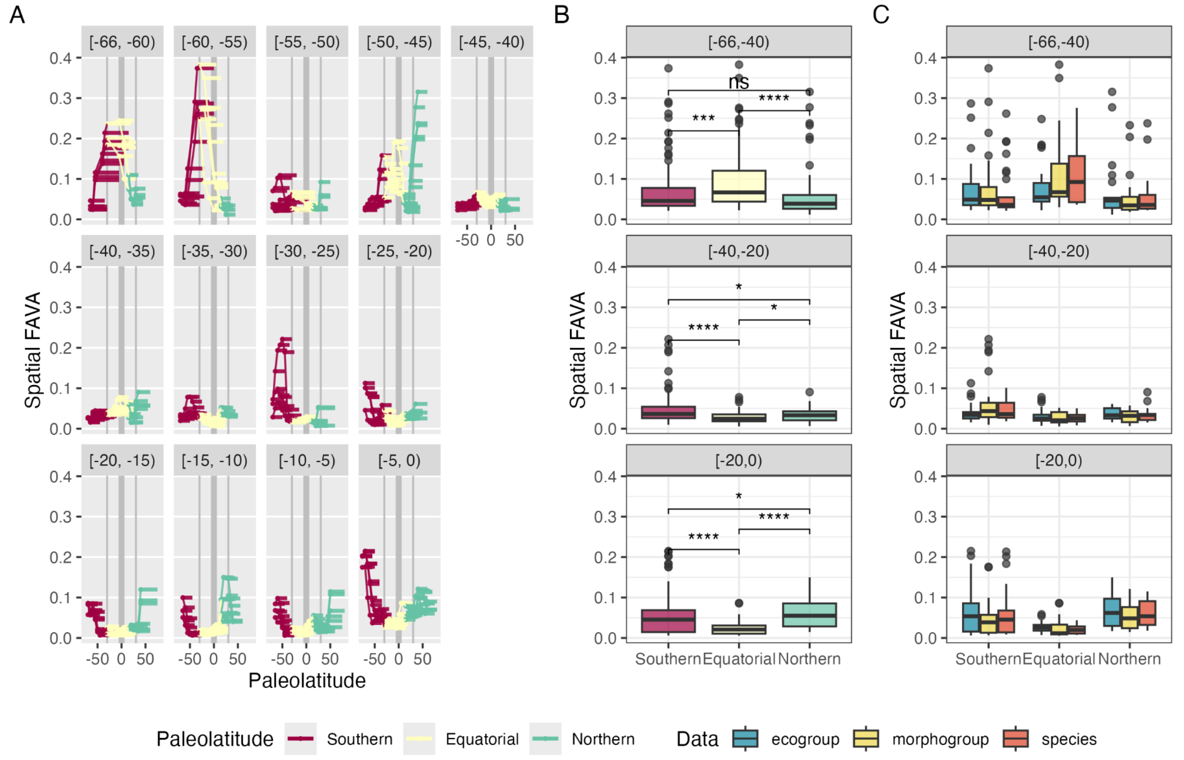


**Figure S8: Latitudinal and temporal gradient in spatial heterogeneity.** (A) Reproduces Figure 2C-D, with data colored by paleolatitude category rather than data type. Thick vertical gay line denotes the equator and thin grey lines denote the boundaries of the equatorial region. Equatorial regions are defined as those whose sliding window is contained entirely between -30 and +30 degrees paleolatitude. (B) Across all data types, the equatorial region is significantly more spatially homogeneous than the southern or northern regions both 0 to 20 Ma and 20 to 40 Ma. However, this pattern reverses 40 to 66 Ma. Asterisks correspond to the magnitude of P-values from Wilcoxon signed-rank tests comparing sFAVA distributions from different paleolatitude regions. These two-sided P-values are as follows:

[-20, 0) Ma: P<10^-6 S:E, P = 0.021 S:N, P < 10^-15 E:N

[-40,-20) Ma: P<10^-6 S:E, P = 0.042 S:N, P = 0.024 E:N

[-66,-40) Ma: P=0.00022 S:E, P = 0.086 S:N, P < 10^-5 E:N

(C) Reproduces Panel B with sFAVA values separated by data type. P-values for each comparison within a data type are as follows:

[-20, 0) Ma, ecogroups: P = 0.013 S:E, P = 0.312 S:N, P < 10^-5 E:N

[-20, 0) Ma, morphogroups: P <10^-6 S:E, P = 0.042 S:N, P =0.024 E:N

[-20, 0) Ma, species: P = 0.0012 S:E, P = 0.174 S:N, P < 10^-7 E:N

[-40,-20) Ma, ecogroups: P = 0.067 S:E, P = 0.959 S:N, P = 0.217 E:N

[-40,-20) Ma, morphogroups: P = 0.0043 S:E, P = 0.081 S:N, P = 0.323 E:N

[-40,-20) Ma, species: P = 0.0019 S:E, P = 0.204 S:N, P = 0.149 E:N

[-66,-40) Ma, ecogroups: P = 0.870 S:E, P = 0.870 S:N, P = 0.340 E:N

[-66,-40) Ma, morphogroups: P = 0.016 S:E, P = 0.052 S:N, P = 0.001  E:N

[-66,-40) Ma, species: P = 0.0013 S:E, P = 0.9558 S:N, P = 0.0025 E:N

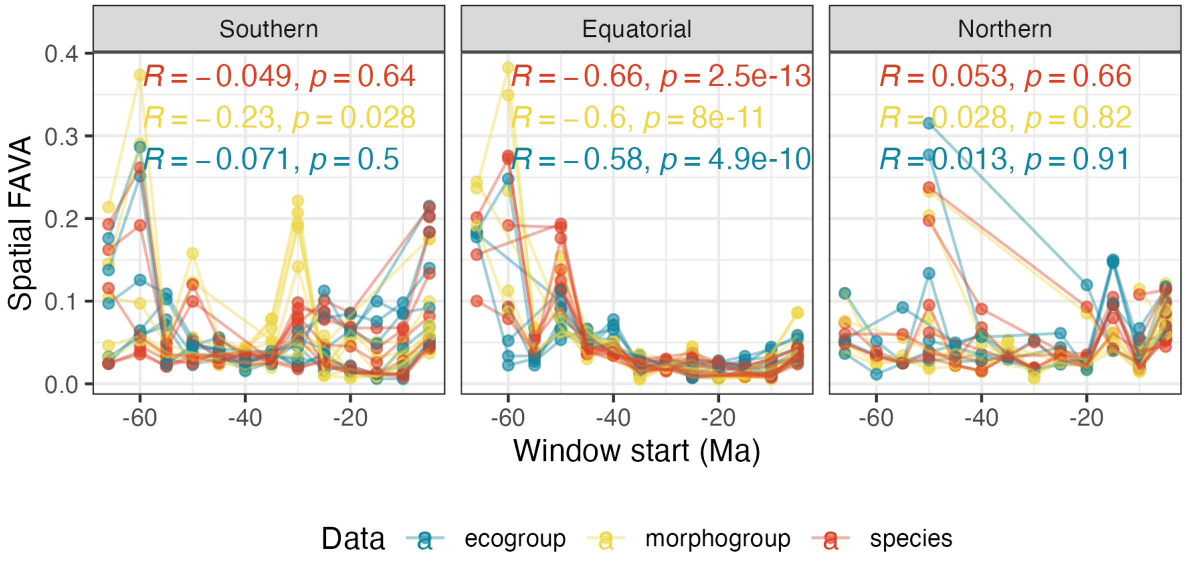


**Figure S9: Temporal trends in spatial heterogeneity depend on latitude.** Each data point corresponds to the start of one window in Figure 2C. Lines connect values of sFAVA in the same paleolatitudinal window through time. Panels present whether these paleolatitudinal windows are southern, equatorial, or northern. Text presents Pearson correlations in each panel for each data set. While there is a slight, negative correlation between window start time and morphogroup sFAVA (R = -0.23, P=0.028), by far the strongest correlations are the negative correlations between time and sFAVA in equatorial regions, regardless of data type. Results of Pearson correlations irrespective of data type are as follows. Southern: R = -0.124, P = 0.038; Equatorial: R = -0.599, P < 10^-15; Northern: R = 0.029, P = 0.678. Results of Pearson correlations irrespective of data type or paleolatitudinal region are: R=-0.265, P < 10^-13.

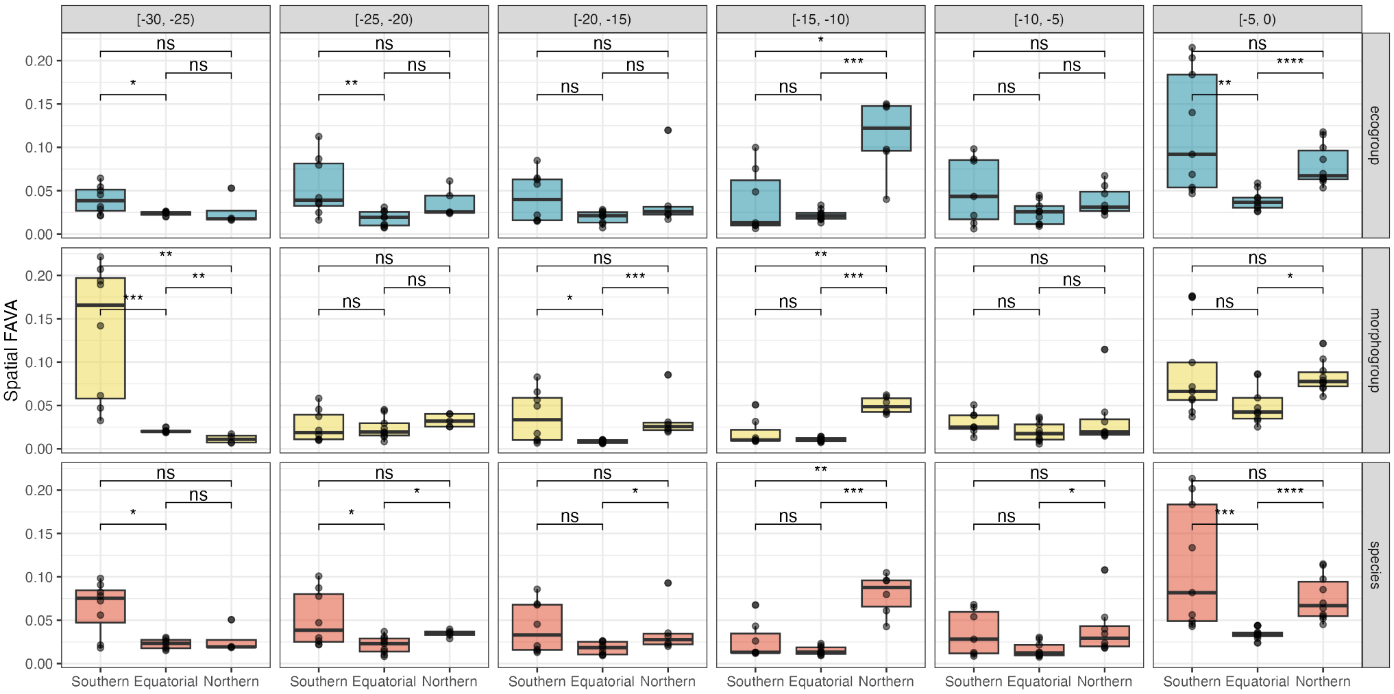


**Figure S10: Latitudinal seesaw of local foraminifera spatial heterogeneity over the past 30 million years.** Each column corresponds to a 5 Myr slice of the last 30 Myrs. Each row presents data from either ecogroups (blue), morphogroups (yellow), or species (red). Each data point represents the spatial FAVA value in one of the 5-degree paleolatitude sliding windows that are depicted in Figure 2C and Figure S8A. "Equatorial" windows are contained within +/- 30 degrees paleolatitude, whereas "southern" windows are south of these boundaries and "northern" windows are to the north. Asterisks present two-sided P-values for Wilcoxon rank sum tests comparing sFAVA values between distinct equatorial regions, in each panel. ns: P > 0.05; *: P <= 0.05; **: P <= 0.01; ***: P <= 0.001; ****: P <= 0.0001.

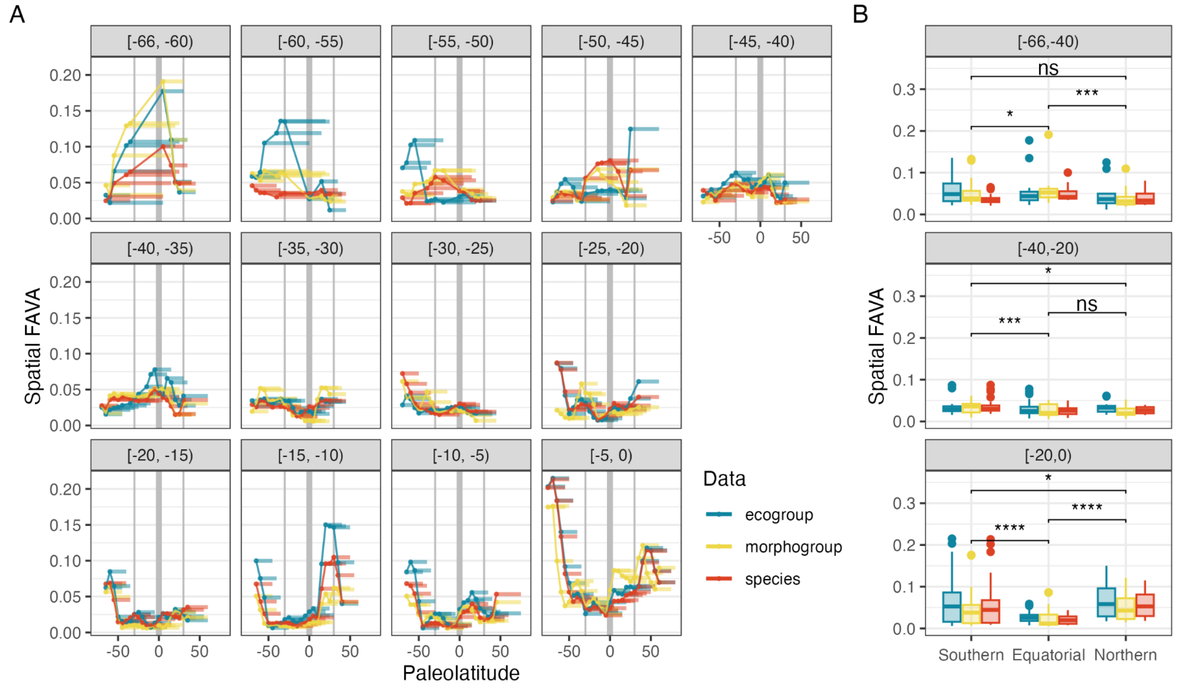


**Figure S11: Trends in spatial FAVA are consistent with a more conservative sampling effort threshold.** Reproduces Figure 2C-D, but with a sampling threshold of 20 rather than 5, eliminating low coverage samples. Consistent with the results presented in Figure 2D, spatial FAVA is still lower in equatorial than in southern (Wilcoxon rank sum test, P<10^-14) or in northern (P<10^-6) regions during the period from 0 to 20 My, and spatial FAVA is still higher in equatorial than in southern (P=0.0118) or northern (P=0.0003) regions.

### Supplementary tables

**Table S1: Quadratic regression results.**

| | statistic | Ecogroup | Morphogroup | Species |
|---|---|---|---|---|
| (Intercept) | estimate | -2.49e-03 | -2.25e-03 | -1.52e-03 |
| *(Intercept)* | *std. error* | *(4.90e-04)* | *(2.90e-04)* | *(2.85e-04)* |
| pal.lat | estimate | -6.96e-06 | 2.81e-06 | -6.79e-09 |
| *pal.lat* | *std. error* | *(1.21e-05)* | *(7.14e-06)* | *(7.02e-06)* |
| I(pal.lat^2) | estimate | 1.07e-06 | 8.42e-07 | 9.84e-07 |
| *I(pal.lat^2)* | *std. error* | *(3.35e-07)* | *(1.98e-07)* | *(1.95e-07)* |
| Num.Obs. | | 21 | 21 | 21 |
| R2 | | 0.505 | 0.570 | 0.675 |
| R2 Adj. | | 0.450 | 0.523 | 0.638 |
| AIC | | -207.6 | -229.7 | -230.4 |
| BIC | | -203.5 | -225.5 | -226.2 |
| Log.Lik. | | 107.817 | 118.858 | 119.195 |
| F | | 9.196 | 11.951 | 18.660 |
| RMSE | | 0.00 | 0.00 | 0.00 |
